# Neural tracking of biological motion rhythms in early infancy: links to caregiver touch-related behaviours and attitudes

**DOI:** 10.64898/2026.05.13.724779

**Authors:** Alicja Brzozowska, Berit Reise, Aneliya Antova, Celine Henning, Stefanie Hoehl

## Abstract

Infant environments are rich in rhythms, many of which are social in nature. These rhythms are proposed to play an important role in early communication and interpersonal synchrony. In this cross-sectional electroencephalography (EEG) study with 3- and 6-month-olds (n=31 and n=30, respectively), we examined whether the infant brain tracks the rhythmicity of locomotion-related biological motion in the visual domain and which experiential factors relate to this ability. We found robust neural tracking of biological motion rhythms at both ages, with no effects of age or orientation (upright or inverted). Additionally, we found that caregiver-reported practice of infant carrying/babywearing and caregiver attitudes toward social touch were linked to infant neural tracking of biological motion rhythms, particularly in the inverted condition. Finally, exploratory analyses revealed a lateralisation effect, whereby the left hemisphere processed rightward (vs. leftward) biological motion rhythms more strongly. Our findings suggest that from early on, the infant brain tracks the rhythmicity of whole-body biological motion. Furthermore, caregiver touch-related practices, particularly infant carrying/babywearing, may play a role in infant neural tracking of socially-relevant rhythms.

## 1. Introduction

Our early environments are abundant in rhythms. From the sound of maternal heartbeat already in the womb, through the rocking movement of the cradle, to the sight of a caregiver shaking a rattle – infants are constantly exposed to rhythmic signals across different modalities (Provasi et al., 2014). Emerging research suggests that the ability of infant brains to track the rhythmicity of perceptual signals, especially those that are socially relevant and informative, may constitute an important mechanism in social and cognitive development (Fiveash et al., 2023; Hoehl et al., 2025; Markova et al., 2019; Provasi et al., 2021). In the present study, we examine infant neural tracking of biological motion, a prominent socially relevant rhythmic signal. We further explore relevant experiential influences by relating this process to infant experience with another type of rhythmic signal – social touch, and particularly the experience of being carried.

The alignment of neural activity with rhythmic perceptual input, referred to as neural tracking or neural entrainment (further used interchangeably), has been observed in the auditory domain as early as in the last semester of human gestation (Saadatmehr et al., 2025). In the first postnatal year, infant brains have been shown to robustly track a wide range of visual and auditory rhythmic stimuli (Kabdebon et al., 2022; Köster et al., 2023). Given that in adults, neural entrainment enhances the processing and encoding of rhythmic stimulus streams (Calderone et al., 2014; Obleser & Kayser, 2019), it emerged as a possible mechanism for learning in early neurocognitive development.

In infancy research, neural entrainment has attracted particular attention in the context of its role in language development (Çetinçelik et al., 2023; Menn et al., 2022; Ní Choisdealbha et al., 2023). Several studies demonstrated that in the first year of life, infants’ brains track quasi-rhythmic signals such as speech (Çetinçelik et al., 2023; Menn et al., 2022; Ortiz Barajas et al., 2021) and singing (Attaheri et al., 2022; Nguyen et al., 2023). Notably, these neural entrainment effects occur not just in the auditory domain, as tracking of the rhythms of syllables and prosodic stress (Menn et al., 2022; Ní Choisdealbha et al., 2023), but also in the visual domain, as tracking of the rhythmic mouth and other facial movements (Ní Choisdealbha et al., 2024), with entrainment being strongest when the stimulus is audio-visual as compared with auditory only (Tan et al., 2022). Importantly, neural entrainment to both visual (Ní Choisdealbha et al., 2024) and auditory (Ní Choisdealbha et al., 2023) features of speech rhythms was found to be a longitudinal predictor of language development. Dynamically aligning brain activity with speech rhythms is thought to amplify the cortical processing of speech and facilitate segmenting the continuous acoustic stream into meaningful units (Obleser & Kayser, 2019). Infants who are better able to track speech rhythms may therefore have an advantage in early language acquisition.

Beyond the auditory domain and speech, entrainment to environmental rhythms has been proposed as a potentially crucial mechanism underlying infants’ ability to dynamically align neural activity and behaviour with other people (Hoehl et al., 2025; Wass et al., 2020). Yet, we are currently lacking empirical evidence for early neural entrainment to non-speech related socially relevant rhythms, especially in the visual domain. In addition to speech-related mouth and other facial movements, there are other examples of rhythmic body movements, the visual processing of which may be relevant for early social and cognitive development. In particular, biological motion associated with others’ locomotion is a ubiquitous, highly rhythmic signal (MacDougall & Moore, 2005; Sydor et al., 2022), and adult brains have been shown to track this rhythmicity in the visual modality (Cracco et al., 2022). Typically studied using point light displays (PLDs) – stimuli representing human movement with a set of moving dots placed on the major joints of the body (Johansson, 1973) – robust processing of biological motion has been demonstrated in the infant brain. Between 5 and 8 months, infants neurally discriminate between upright and inverted (Marshall & Shipley, 2009), scrambled and intact (Lisboa et al., 2020), and biomechanically possible and impossible (Reid et al., 2008) biological motion. However, to the best of our knowledge, no study to date has examined whether infant brains track the rhythmicity of biological motion. In analogy to speech tracking, biological motion tracking could facilitate neural processing of others’ actions and aid, for instance, action segmentation or the detection of synchronous movement. If this is the case, individual differences in this ability may have cascading effects on later social-cognitive abilities, argued to be scaffolded by early biological motion processing, such as imitation (Happé et al., 2017) and empathy (Miller & Saygin, 2013; Rice et al., 2016).

Further, despite strong premises to hypothesise that biological motion processing is susceptible to learning (Hirai & Senju, 2020; Pavlova, 2012), there is hardly any research on what factors shape this learning process in early development. Initial evidence points to the influence of the infant’s own action experience: artificial introduction of walking experience at 10 weeks was shown to enhance processing of upright, as compared with inverted biological motion (Reid et al., 2019). Indeed, adolescents with cerebral palsy who had experienced motor disabilities from birth show impairments in gesture meaning recognition from biological motion (Maryniak & Foryś – Basiejko, 2025), further implicating one’s own motor experience as an important developmental factor. Yet, pre-walking infants show robust predictive tracking of stepping actions, suggesting that one’s own experience is not necessary for sophisticated processing of action kinematics (De Klerk et al., 2016).

In line with this observation, it has also been argued that visual experience and perceptual learning may be responsible for gradual improvements in biological motion processing (Pavlova, 2012), although evidence supporting this notion comes exclusively from studies with adults trained to develop expertise in arbitrary forms of non-human biological motion (Grossman et al., 2004; Jastorff et al., 2009). Interestingly, individuals who had experienced a temporary period of congenital visual deprivation caused by congenital cataracts show impairments in face processing (Robbins et al., 2010), but do not differ from matched controls in their processing of biological motion (Bottari et al., 2015), suggesting potential multimodal contributions to biological motion processing proficiency.

We argue that there are compelling reasons to examine caregiver touch and proximity as factors that may relate to biological motion processing in early development. In particular, the practice of infant carrying/babywearing may be especially relevant for the processing of rhythmic aspects of biological motion. While being carried, infants experience multimodal – auditory, vestibular, and proprioceptive – rhythmic stimulation at the frequency of their caregiver’s locomotion (Rocha et al., 2021b, 2021a). Through cross-modal effects on rhythm processing (Phillips-Silver & Trainor, 2005; Russo et al., 2024), this stimulation is likely to affect infant processing of closely related rhythms in the visual modality. As humans tend to synchronise while walking next to each other (Chambers et al., 2019; Hoch et al., 2021), it is probable that infants would have plenty of such correlated multimodal experience, potentially leading to enhanced neural tracking of biological motion rhythms in infants who are carried a lot.

Additionally, caregiver touch and proximity have been proposed to act as a social communicative cue, enhancing the salience of social signals (Addabbo et al., 2025; Akhtar & Gernsbacher, 2008; Carozza & Leong, 2021). Previous research showed that while experience of affective touch appears to be unrelated to overt measures of social attention in the first year of life, such as looking towards social as compared with non-social stimuli (Brzozowska et al., 2022; Della Longa et al., 2019; Nava et al., 2020), it may affect covert aspects of social cognition. For instance, 4-month-olds remembered faces with averted gaze better when their presentation was accompanied by stroking by hand as compared with brush tapping or no touch, despite a lack of looking time differences between the three conditions (Della Longa et al., 2019). Further, proximity and affective touch are linked to measures of neural (but not physiological) synchrony between 4-6-month-olds and their caregivers (Nguyen et al., 2021). Finally, in older children, at around 5 years, touch received from caregivers during a free play interaction is positively correlated with resting activity within the so-called “social brain” network (Brauer et al., 2016), which includes the superior temporal sulcus – a region known to be involved in biological motion processing (Pelphrey & Morris, 2006). In sum, prior research suggests that there may be a (potentially causal) link between caregiver touch and infant processing of socially relevant stimuli – including biological motion – and that this link may be most evident in measures capturing covert aspects of social cognition, such as neural activity measures.

The aims of the present study were as follows: first, we aimed to examine whether infant brains track the rhythmicity of biological motion in the first months of life. As we were interested in developmental changes in biological motion processing, we recruited two age groups: 3- and 6-month-olds. The inclusion of the younger age group was motivated by the apparent research gap in neuroimaging studies on infant processing of biological motion, which thus far have focused on the 5 – 8 months age range (Lisboa et al., 2020; Marshall & Shipley, 2009; Reid et al., 2006, 2008); the older group was included as a better studied population, known to exhibit robust neural processing of different aspects of biological motion. Based on theoretical accounts arguing for improvements in biological motion processing across development (Pavlova, 2012), we hypothesised that we would observe stronger neural tracking of upright, as compared with inverted, biological motion PLDs in both age groups, but that this effect would be stronger in the older group. Second, we aimed to examine the possible links between infant neural tracking of biological motion rhythms and caregiver-reported touch related behaviours and attitudes. Specifically, we hypothesised that, due to its possible effects on rhythmic processing, caregiver-reported practice of infant carrying and babywearing will be positively related to infant neural tracking of biological motion. Further, through more general effects on social information processing, caregiver-reported touch behaviours and attitudes will also be positively related to infant neural tracking of biological motion. For these two hypotheses, an interaction with the orientation of PLDs was expected, such that the effects would be more pronounced for the upright PLDs.

To test our hypotheses, we recorded electrical brain activity from infants using electroencephalography (EEG) while they were viewing PLDs moving at a rhythmic pace, using an adapted paradigm previously applied with adult participants (Cracco et al., 2022). Additionally, we collected caregiver-report measures of infant carrying, as well as infant-directed touching behaviours (Brzozowska et al., 2021; Koukounari et al., 2015), and caregivers’ attitudes and affect associated with social touch (Wilhelm et al., 2001).

## 2. Materials and methods

This study has been preregistered, the preregistration document is available on the platform AsPredicted: https://aspredicted.org/ev22y6.pdf. Minor deviations from the preregistration are detailed in section S1 in Supplementary Materials. The analysis scripts and data supporting the results of the study are openly available at https://github.com/alicja444/BioMot and https://doi.org/10.17605/OSF.IO/THFYE, respectively. The study received approval of the Ethics Committee of the University of Vienna (ref. 01128).

### 2.1. Participants

The infants who participated in the study were full-term, typically developing 3-month-olds (n = 41, mean age = 107 days; 21 female and 20 male) and 6-month-olds (n = 41, mean age = 200 days; 19 female and 22 male). Ten 3-month-olds and eleven 6-month-olds were tested but not included in the analyses due to fussiness and/or insufficient data quality (see Supplementary Materials, section S2 for details on reasons for data missingness), yielding a final sample of 31 3-month-olds and 30 6-month-olds. The testing took place in the Vienna Children’s Studies lab (Faculty of Psychology, University of Vienna, Austria). Families recruited in the study were local to Vienna, i.e., the sample was drawn from an urban environment. In the 3-month-olds group, 75% of families were exclusively German-monolingual, while in 25% of families infants were exposed to at least two languages; in the 6-months-old group, these percentages were 68% and 29%, respectively (3% missing data). We assessed participants’ communities of descent with an open-ended question about the cultural identification of the (self-identified) primary caregiver and, if applicable, the secondary caregiver. Such approach is considered appropriate in the Austrian cultural context (Vietze et al., 2022). Sixty-five percent of primary caregivers reported exclusively Austrian cultural identification, 10% both Austrian and other cultural identification, 20% reported exclusively non-Austrian cultural identification(s), while 5% broadly described their cultural identification as European. Among secondary caregivers, 58% reported exclusively Austrian cultural identification, 10% both Austrian and other cultural identification, 25% reported exclusively non-Austrian cultural identification(s), and 4% a broad European identification; for 3% of secondary caregivers this information was missing. Among primary caregivers, 11% reported their highest level of education as upper secondary, while 89% reported tertiary; among secondary caregivers these percentages were 24% and 74%, respectively (2% missing data).

### 2.2. Stimuli and procedure

#### 2.2.1. Biological motion task

The biological motion task used in the present study was adapted from Cracco et al. (2022). Infants were presented with video clips of point light walkers in 4 conditions, varying in orientation (upright/inverted) and direction (left/right). All point light walkers were moving at a frequency of 2.4 Hz, i.e., one step every ∼417 milliseconds. At the beginning of each trial, an attention grabber (cartoon drawing of a cloud) was presented in the middle of the screen for a random duration between 500 and 750 milliseconds. Next, a video of one of the four conditions was shown for 10 seconds, accompanied by child-friendly music matching in tempo (144 BPM), with four different song snippets counterbalanced across conditions. There were 32 trials in total, each randomly assigned one of the four experimental conditions, (totalling in 8 trials per condition) with no more than one repetition of a condition in a row. In case of the infant’s continued interest in the displays, further trials (following the same pattern) were presented until the infant became fussy. The experimental task is shown in Figure 1.

**Figure 1.**
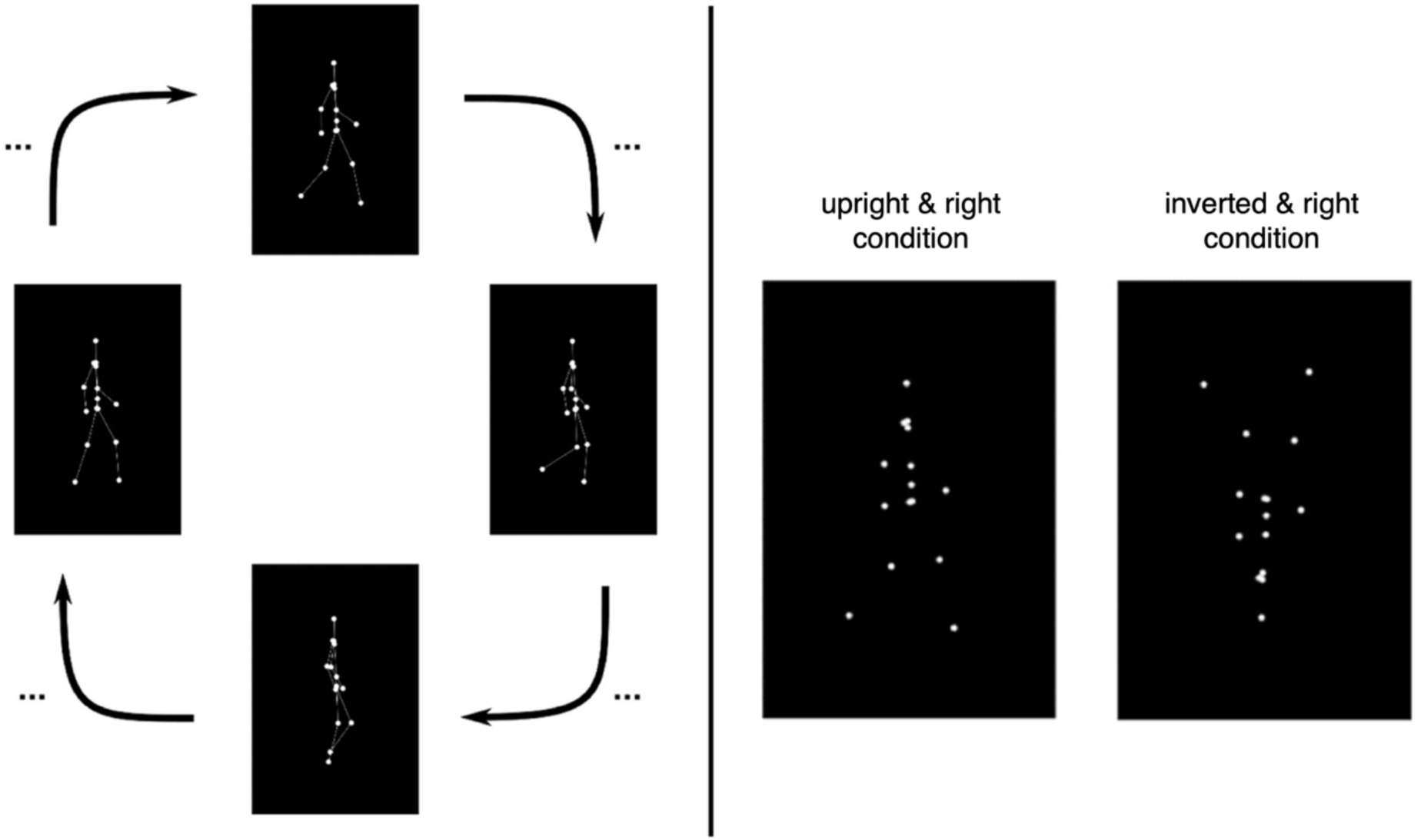
Biological motion task, adapted from Cracco et al. (2022); licensed under CC-BY. Left: example frames showing movement progression of the point light walker (lines between dots not shown in the actual experiment); right: example frames from the upright and right and inverted and right conditions (additional conditions not depicted: upright and left, inverted and left).

#### 2.2.2. Parent-Infant Caregiving Touch Scale (PICTS)

The Parent-Infant Caregiving Touch Scale (PICTS) is a parent-report instrument assessing touch-related behaviours of caregivers towards their infants (Koukounari et al., 2015). The questionnaire consists of simple items such as “I cuddle my baby” or “I stroke my baby’s tummy”, and the response is recorded on a Likert scale ranging from 1 (never) to 5 (a lot). We used an adapted version translated into German, which was previously shown to relate to parental touching behaviors as observed during free play in the lab (Brzozowska et al., 2021).

The adapted version included an extra item about infant carrying (see below). The total score was calculated as a sum of responses to individual items, with reverse-scored response to the item “I leave my baby to lie down”.

#### 2.2.3. Infant Carrying

Infant carrying was assessed with a single item from the adapted PICTS questionnaire: “I carry my baby in a sling, or similar” (German: *Ich trage mein Baby in einem Tragetuch o.ä.*). The score ranged from 1 (never) to 5 (a lot).

#### 2.2.4. Social Touch Questionnaire

The Social Touch Questionnaire (STQ) is a questionnaire measuring attitudes towards situations involving social touch (Wilhelm et al., 2001). It consists of items such as “I feel uncomfortable when someone I don’t know very well hugs me” or “I generally seek physical contact with others”, with the response on a Likert scale ranging from 0 (not at all) to 4 (extremely). Although the questionnaire is designed to measure general touch-related attitudes and affect, it has previously been used specifically in the context of caregiver-infant touch (Aguirre et al., 2019; Fairhurst et al., 2014). Our adapted version was translated into German and did not include three items present in the original questionnaire, as they were deemed either not applicable to our study participants (“I’d feel uncomfortable if a professor touched me on the shoulder in public”) or associated with romantic, intimate touch (“I like being caressed in intimate situations” and “I feel disgusted when I see public displays of intimate affection”). This adapted version of STQ was found to relate to caregiver infant-directed touch as observed in the lab (Brzozowska et al., 2021). A recently published German version of the STQ, very similar to ours but containing all original items, was reported to have good psychometric properties (Lapp & Croy, 2021). As the questionnaire was originally developed to focus on anxiety and embarrassment associated with social touch in the context of social anxiety (Wilhelm et al., 2001), in its original version higher scores represent more negative attitudes and affect toward social touch. In the present study, we reverse-scored the questionnaire so that higher scores represent more positive attitudes and affect toward social touch; for clarity, we further refer to this score as STQ-r.

#### 2.2.5. Procedure

Testing took place in a room adapted for developmental EEG data collection. First, the study procedure was explained to the caregiver and written consent was collected. Next, an EEG cap was placed on the infant’s head and gel was added to the electrodes. The infant was then seated on the caregiver’s lap, approximately 60 cm from the screen. The biological motion task was then displayed on an Iiyama G-Master GB2560HSU screen with a 144 Hz refresh rate, and the accompanying audio was presented using Logitech Multimedia Z200 speakers. After the biological motion task was completed and the EEG gel was washed off the infant’s head, the caregiver was asked to fill out the two short questionnaires (PICTS, STQ) and a brief socio-demographic form. At the end of the visit, the family’s travel costs to the lab were reimbursed and the infant received an age-appropriate toy and a certificate of participation. The entire visit in the lab lasted about an hour.

### 2.3. Data collection, preprocessing, and analysis

#### 2.3.1. Looking behavior

Infant looking behavior towards the screen was recorded with two Axis P1365 Mk II cameras at 25 frames per second. These videos were later used for off-line coding of infant gaze (on/off the screen) with the use of the Interact software (Mangold, 2018).

#### 2.3.2. EEG data collection

We used EEG caps (EasyCap GmbH) in combination with an active 32 Ag/AgCl electrode set (ActiCap by Brain Products GmbH) and conductive gel in data collection. The impedances were checked right before the start of data collection to ensure that they were below 10 kΩ, whenever possible. The placement of the electrodes was an adapted 10–20 layout system, with the online reference electrode positioned behind the left ear (TP9), and the ground electrode in the front (Fp1). For EEG data acquisition we used a BrainAmp DC amplifier (Brain Products GmbH), with a sampling frequency of 500 Hz. A Photo Sensor (Brain Products GmbH) was used to generate precise triggers of trial onsets and offsets. We used BrainVision Recorder software (BrainVision Recorder, Vers. 1.24.001, Brain Products GmbH, Gilching, Germany) to record the EEG signal. Triggers from the Photo Sensor, the experimental script, and the EEG and video recording software were synchronised with a TriggerBox device (Brain Products GmbH).

#### 2.3.3. EEG data preprocessing

The EEG data was preprocessed with the use of the standardized HAPPILEE pipeline for lower density recordings (Lopez et al., 2022) in combination with custom MATLAB (The MathWorks Inc, 2022) scripts, and functions from the EEGLAB toolbox (Delorme & Makeig, 2004). Firstly, event triggers were updated so that they corresponded to the latencies of the Photo Sensor triggers, ensuring precise timing. Line noise processing was performed with the default CleanLine method of the HAPPILEE pipeline. The continuous signal was then bandpass-filtered with a zero-phase Hamming-windowed sinc FIR filter with cut-off frequencies 1 Hz and 100 Hz. Wavelet thresholding was used for artifact correction. Data within trials was then segmented into 1.668-second epochs (duration of 4 full cycles of the 2.4 Hz oscillation).

Channels identified by the HAPPILEE pipeline as artifact-contaminated were interpolated within epochs using spherical spline interpolation (no more than 3 channels per epoch, otherwise the epoch was rejected). Epochs were then automatically rejected if the amplitude in any of the channels within the region of interest (parieto-occipital electrodes: P7, P3, Pz, P4, P8, PO9, O1, Oz, O2, PO10) was below/above +/- 150 μV. The performance of the HAPPILEE pipeline was inspected in the automatically-generated quality assessment file, and participants for whom cross-correlation values across all frequencies before and after wavelet thresholding were below 0.1 (indicating major signal change) were excluded from further analyses (Monachino et al., 2022). The data was then re-referenced to average reference and visually inspected for quality. Additional individual epochs were then rejected manually if significant quality issues were identified. Epochs where infants’ gaze was coded as directed away from the screen (for any duration of the epoch) were excluded from further analyses.

#### 2.3.4. EEG data analysis

The EEG data was analysed using custom MATLAB (The MathWorks Inc, 2022) scripts and functions from the EEGLAB toolbox (Delorme & Makeig, 2004). As preregistered, and following relevant prior work (Cracco et al., 2022; Reid et al., 2006) our analyses focused on the parieto-occipital sites (electrodes P7, P3, Pz, P4, P8, PO9, O1, Oz, O2, and PO10). Epochs were averaged across conditions and a fast Fourier transform was applied to zero-padded averaged signals, yielding a spectral resolution of 0.4 Hz. Baseline-corrected amplitude spectra were computed by subtracting the average of amplitudes at two (not immediately) neighbouring frequency bins, +/- 0.8 Hz from each frequency, similarly to Cracco et al. (2022). Baseline-subtracted amplitudes at the three (preregistered) frequencies of interest: the fundamental frequency, 2.4 Hz, and its two harmonics, 4.8 Hz, and 7.2 Hz were then extracted for all participants, separately for each condition and hemisphere (left/right), as previous research suggested that the response of interest may be lateralized (Cracco et al., 2022; Reid et al., 2006). Further analyses were performed on these extracted baseline-subtracted amplitudes.

### 2.4. Analytic strategy

The analytic plan was as follows: First, we explored infant looking durations to stimuli depending on the stimuli orientation, direction, and infants’ age (for all participants, including those who did not contribute EEG data). Then, we examined correlations between touch related-measures. Next, the presence of neural tracking was verified by testing the baseline-corrected amplitudes averaged across conditions at the three harmonics of interest (2.4 Hz, 4.8 Hz, and 7.2 Hz), separately for each age group, against the value of 0. Values significantly greater than 0 indicated the presence of neural tracking and validated the inclusion of a given harmonic in further analyses. The next step was to examine the preregistered hypotheses: H1: Neural entrainment to biological motion is observable at 3 months and at 6 months, but stronger at 6 months, and H2: Levels of infant carrying and other infant-directed touching behaviours reported by the primary caregiver(s) will be positively correlated with neural entrainment to biological motion at 3 and at 6 months. The hypotheses were examined with the use of mixed-effect models (small deviations from preregistered procedures are outlined in Supplementary Materials, section S1). Finally, an exploratory analysis that included additional factors such as the number of included epochs, direction of movement of the point light walkers, and hemisphere in which neural tracking was measured was conducted. This was done through gradually adding fixed effects of interest to the original, preregistered model and selecting the best model using Likelihood Ratio Tests for model comparisons (Muradoglu et al., 2023).

## 3. Results

### 3.1. Descriptive statistics, looking behaviour and relations between touch measures

At 3 months, 75% of participants contributed valid EEG data while at 6 months, this proportion was 73%. Detailed information regarding reasons for missingness is reported in Supplementary Materials (section S2). Descriptive statistics of the main variables in the study are shown in Table 1.

**Table 1.** Descriptive statistics of the neural tracking data and caregiver-report touch-related measures. Note that the baseline-subtracted amplitudes were summed across the two harmonics (2.4 Hz and 4.8 Hz).

|  | 3 months |  |  |  |  | 6 months |  |  |  |  |
| --- | --- | --- | --- | --- | --- | --- | --- | --- | --- | --- |
|  | n | Mean | SD | Minimum | Maximum | n | Mean | SD | Minimum | Maximum |
| **Amplitude [μV] in the upright condition** | 31 | 0.26 | 0.28 | -0.33 | 0.99 | 30 | 0.18 | 0.29 | -0.54 | 0.68 |
| **Amplitude [μV] in the inverted condition** | 31 | 0.30 | 0.32 | -0.31 | 1.09 | 30 | 0.31 | 0.40 | -0.49 | 1.30 |
| **PICTS** | 25 | 53 | 5 | 44 | 62 | 30 | 53 | 5 | 43 | 61 |
| **Infant carrying** | 25 | 4 | 1 | 2 | 5 | 30 | 4 | 1 | 2 | 5 |
| **STQ-r** | 25 | 40 | 9 | 23 | 56 | 30 | 44 | 8 | 28 | 56 |

Further, we explored how long infants looked at the point-light walkers during the 10-second-long trials. At 3 months, infants looked on average 8629 ms (Mdn = 9626 ms, SD = 2521 ms) at the stimuli, 8679 ms in the inverted and 8580 ms in the upright condition. At 6 months, infants’ mean looking time was 8275 ms (Mdn = 9136 ms, SD = 2216 ms). On average, they looked 8235 ms at the stimuli in the inverted and 8315 ms in the upright condition. We then fitted a mixed-effects model predicting looking duration with main effects of Direction, Orientation and Age, an interaction term between Orientation and Direction, and a random effect of participant (intercept only). As per recommendation for infant looking data (Csibra et al., 2016), we log-transformed looking durations.

We found no significant effects. The main effects of Orientation (β = 0.02, SE = 0.02, t(2260) = 0.87, p = 0.38), Direction (β = 0.02, SE = 0.02, t(2260) =1.17, p = 0.24), and their interaction (β = 0.00, SE = 0.02, t(2233) = 0.26, p = 0.8) were not significant. Neither was the main effect of age (β = 0.1, SE = 0.07, t(72) = 1.49, p = 0.14).

Next, we examined associations between the three touch-related variables of interest (PICTS, Infant Carrying, and STQ-r) and found that while PICTS and STQ-r were moderately correlated (r(69) = 0.34, p =.003), neither PICTS (r(69) = 0.22, p =.07), nor STQ-r (r(69) = - 0.19, p =.12) correlated with Infant Carrying.

### 3.2. Verifying the presence of neural tracking

First, we verified the presence of neural tracking by testing the response at the fundamental frequency (2.4 Hz) and its two harmonics (4.8 Hz and 7.2 Hz) against the value of 0 in both age groups.

At 3 months, the 2.4 Hz response was significant both in the upright (t(30) = 3.78, p <.001) and inverted (t(30) = 4.28, p <.001) condition. The 4.8 Hz response was also significant in the upright (t(30) = 4.24, p <.001) and inverted (t(30) = 4.54, p <.001) condition. However, the 7.2 Hz harmonic was not significant in either the upright (t(30) =-1.38, p =.18) or inverted condition (t(30) =-0.45, p =.66).

At 6 months, the 2.4 Hz response was significant in the upright (t(29) = 2.33, p =.03) and inverted (t(29) = 3.64, p =.001) condition. The 4.8 Hz harmonic was significant as well, in both the upright (t(29) = 2.54, p =.02) and inverted (t(29) = 2.70, p =.02) conditions. Similarly to the 3-month-olds, 6-month-olds did not show a significant response at 7.2 Hz at either the upright (t(29) = 1.14, p =.26) or inverted (t(29) = 0.93, p =.36) conditions. The lack of significance of the 7.2 Hz harmonic in both age groups led us to exclude it from further analyses. Consequently, all the analyses that follow include only the 2.4 Hz and 4.8 Hz responses. The amplitude spectra for the 3- and 6- month-olds are shown in Figure 2.

**Figure 2.**
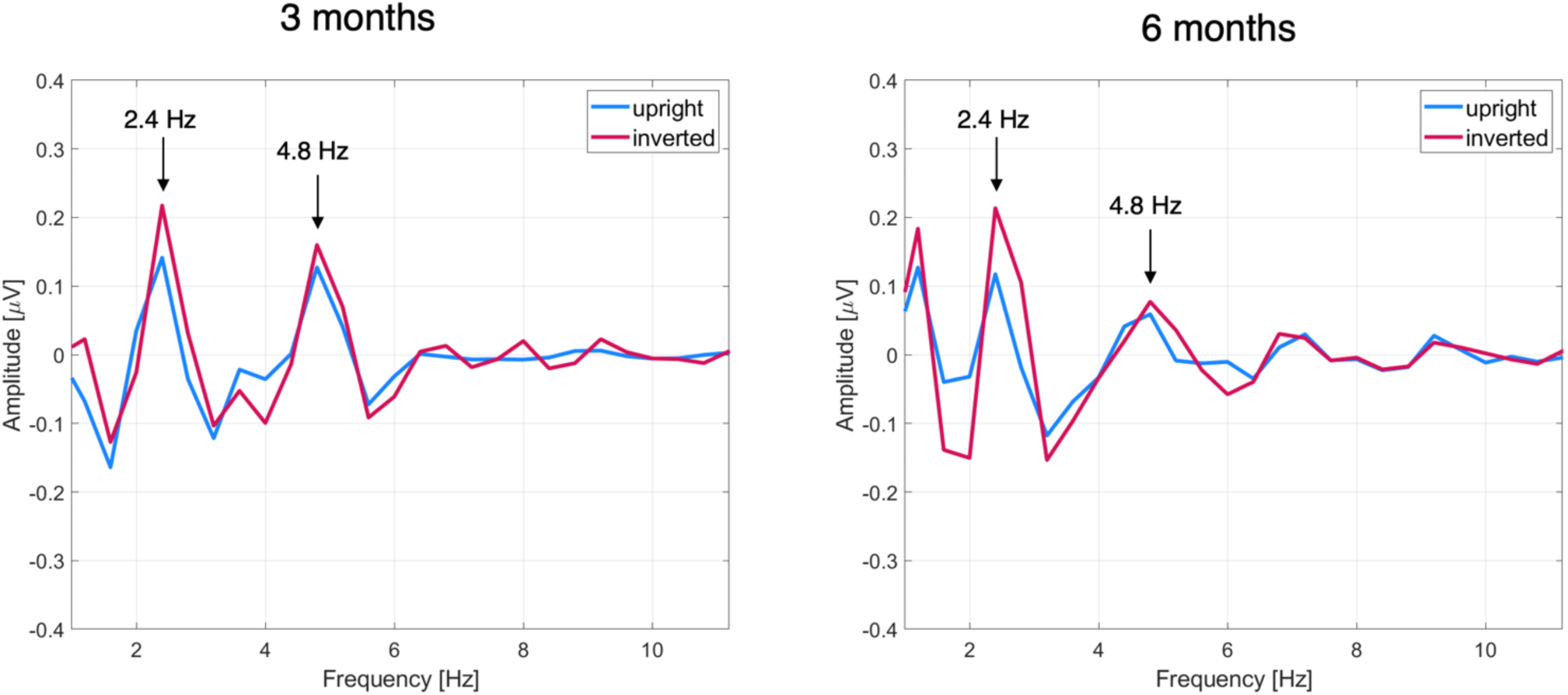
Baseline-subtracted amplitude spectra averaged across participants at 3 months (left) and 6 months (right). Additionally, we checked whether the number of available data segments differed between Orientation conditions by fitting a mixed-effects model predicting the number of available segments with a fixed effect of Orientation (upright/inverted) and a random effect of participant (intercept only). The effect of Orientation was not significant (β =-0.03, SE = 0.02, t(914) =-1.34, p =.18), meaning that there were no significant differences in the number of available data segments between the upright and inverted conditions.

### Preregistered analysis: Does neural tracking of biological motion improve with age?

To test the preregistered hypothesis that neural tracking of upright, but not inverted, biological motion improves with age, we fitted a mixed-effects model predicting baseline-subtracted amplitudes at the two harmonics of interest with Orientation (upright/inverted), Age (3 months/6 months), and an interaction between Orientation and Age as fixed effects, as well as a random effect of participant (intercept only). We found no evidence for a main effect of Orientation (β = - 0.05, SE = 0.09, t(913) =-0.60, p =.55), a main effect of Age (β = 0.02, SE = 0.11, t(137) = 0.16, p =.88), or an interaction between Orientation and Age (β = - 0.13, SE = 0.12, t(913) =-1.02, p =.31). The intercept was β = 0.05, SE = 0.07, t(137) = 0.67, p =.51.

### Preregistered analysis: Is infant neural tracking of biological motion rhythms linked to caregiver-reported levels of infant carrying, other touch related behaviours, and attitudes toward social touch?

To test the preregistered hypothesis that levels of infant carrying, other touch-related behaviours, and caregiver attitudes toward social touch as reported by the primary caregiver will positively predict infant neural tracking of biological motion, we fitted a mixed-effects model predicting baseline-subtracted amplitudes with main effects of Orientation, PICTS score, Infant Carrying score, and STQ-r score, and three interaction terms: between Orientation and PICTS, between Orientation and Infant Carrying and between Orientation and STQ-r, as well as a random effect of participant (intercept only). We found that the main effects of Orientation (β =-0.11, SE = 0.07, t(134) =-1.69, p =.09), PICTS (β = - 0.10, SE = 0.06, t(134) =-1.82, p =.07), and their interaction (β = 0.07, SE = 0.07, t(821) = 0.98, p = 0.33) were not significant. However, we found evidence for a main effect of Infant Carrying (β = 0.16, SE = 0.06, t(134) = 2.93, p =.003), a main effect of STQ-r (β = 0.14, SE = 0.06, t(134) = 2.36, p =.02) and an interaction between Orientation and Infant Carrying (β =-0.15, SE = 0.07, t(821) =-2.14, p =.03) as well as between Orientation and STQ-r (β = - 0.23, SE = 0.07, t(821) =-3.10, p =.002).

The intercept was β = 0.06, SE = 0.05, t(134) = 1.08, p =.28. The significant interaction effects from this model are shown in Figure 3.

**Figure 3.**
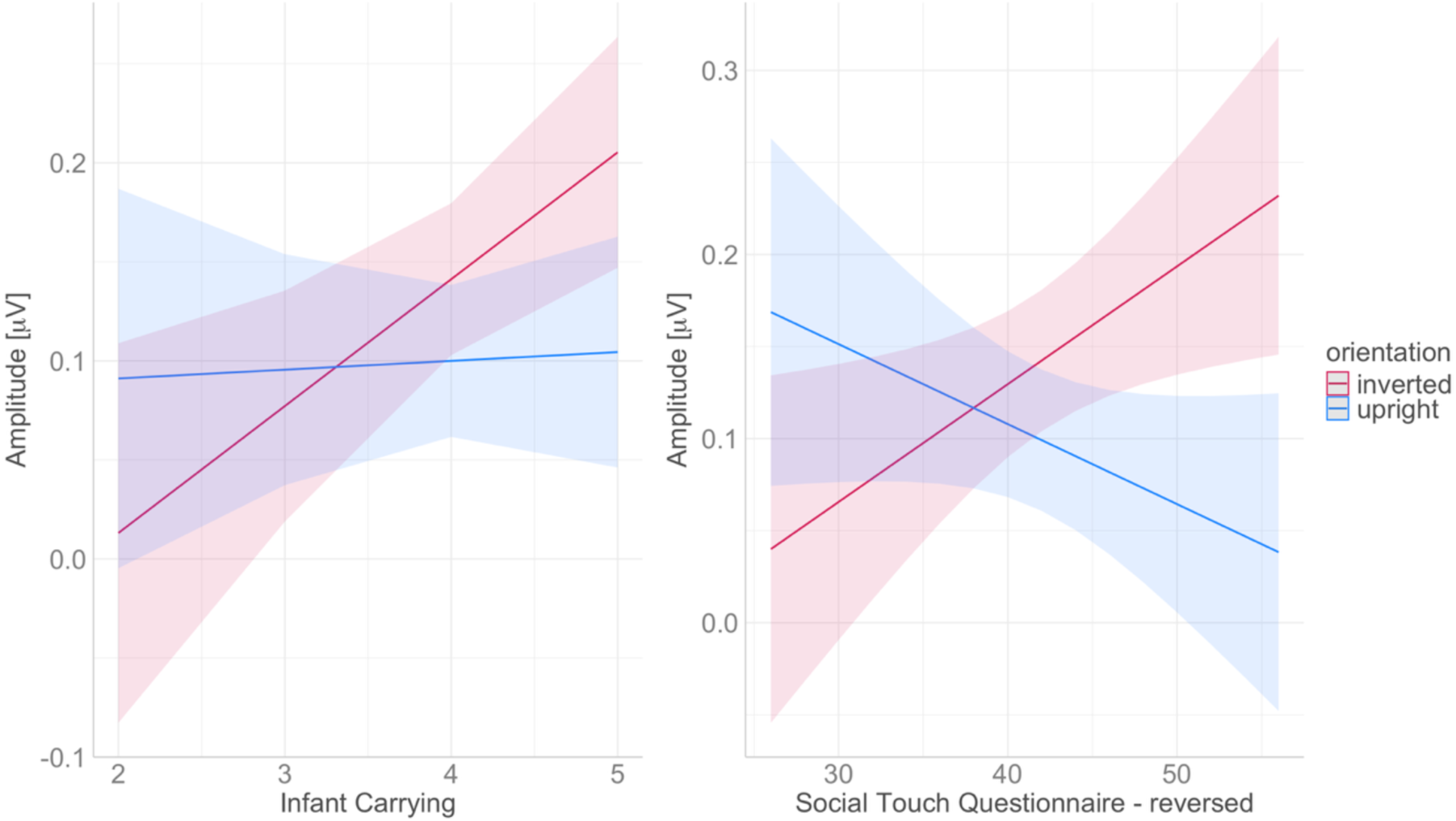
Interaction effects between Infant Carrying and Orientation (left) and Social Touch Questionnaire – Reversed and Orientation (right) on baseline-subtracted amplitudes.

The interaction effects suggest that the main effects showing a positive link of Infant Carrying and STQ-r with baseline-subtracted amplitudes were driven by the inverted condition of Orientation. At high levels of Infant Carrying and STQ-r, infants’ neural tracking of upright and inverted biological motion appears to differ, being stronger in the inverted condition.

### Exploratory analysis: examining additional factors in neural tracking of rhythmic biological motion

Next, we explored additional factors that may affect the strength of neural tracking of rhythmic biological motion in infants, including Direction (left/right), Hemisphere in which the neural response is measured (left/right parieto-occipital sites), and Number of data segments available. We also considered potential interactions of the effects with Age. Starting with our preregistered model described in the previous section, and gradually adding fixed effects of interest, we selected the best model using Likelihood Ratio Tests for model comparisons. An overview of all models considered in the model selection process can be found in Table 2.

**Table 2.**
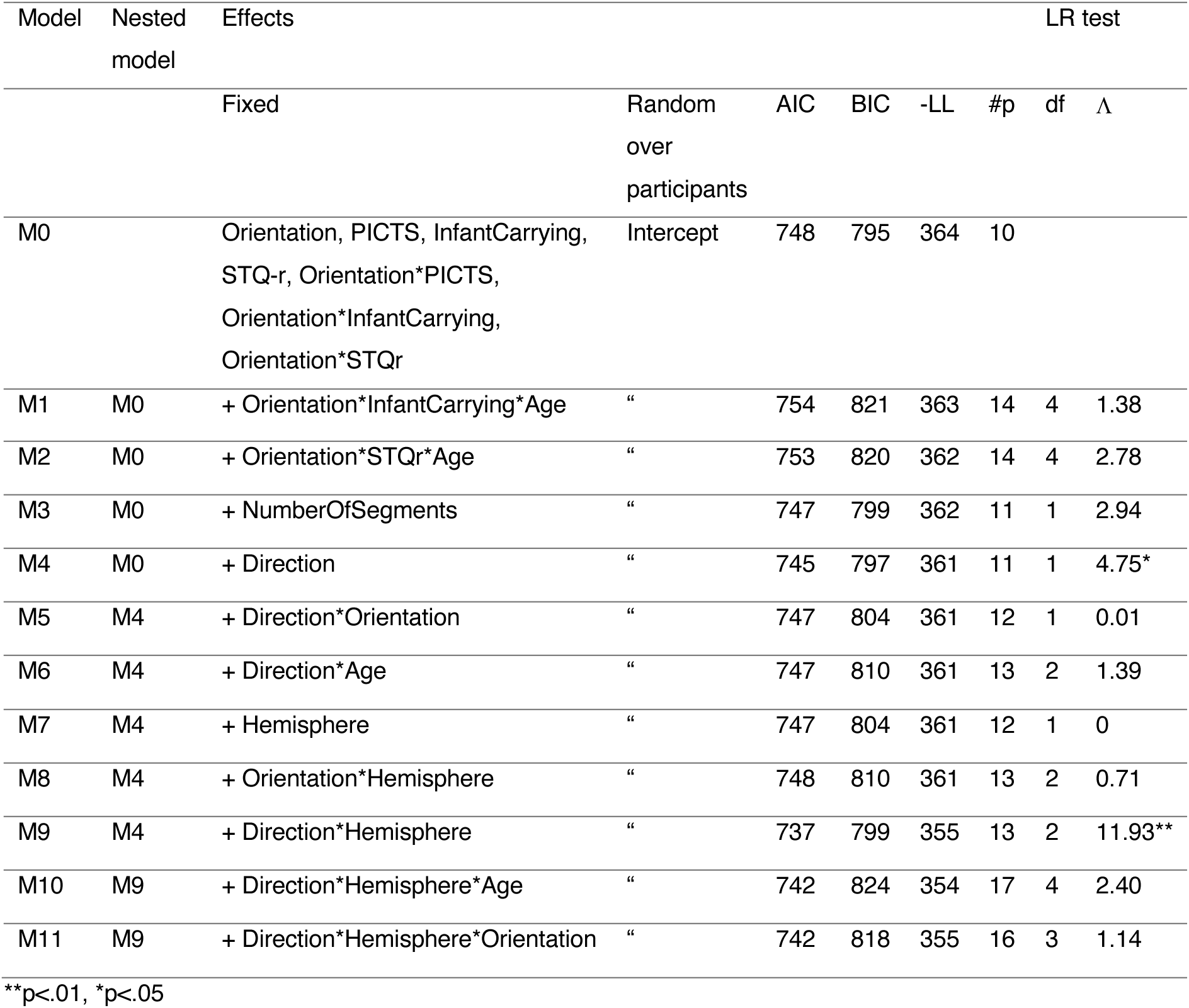
Overview of the estimated mixed-effects models.

The best model (M9), in addition to the parameters present in the original model (random effect of participant - intercept only, fixed effects of Orientation, PICTS, Infant Carrying, STQ-r, as well as interactions between PICTS and Orientation, Infant Carrying and Orientation, and STQ-r and Orientation), included fixed main effects of Direction and Hemisphere, and an interaction between them. Details of the estimated fixed effects in the best model are included in Table 3.

**Table 3.**
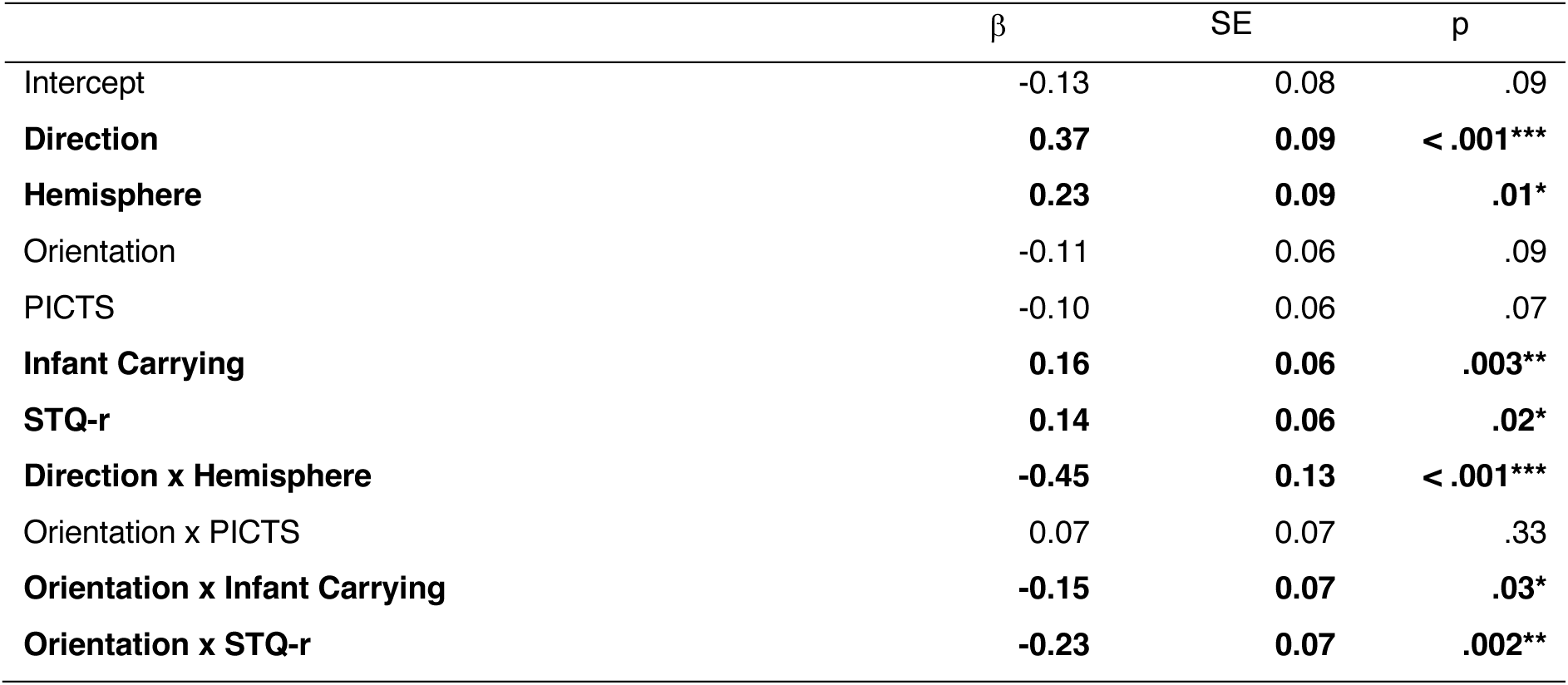
Estimates of fixed effects in the best-fitting model (M9).

| | $\beta$ | SE | p |
| --- | --- | --- | --- |
| Intercept | -0.13 | 0.08 | .09 |
| **Direction** | **0.37** | **0.09** | **< .001\*\*\*** |
| **Hemisphere** | **0.23** | **0.09** | **.01\*** |
| Orientation | -0.11 | 0.06 | .09 |
| PICTS | -0.10 | 0.06 | .07 |
| **Infant Carrying** | **0.16** | **0.06** | **.003\*\*** |
| **STQ-r** | **0.14** | **0.06** | **.02\*** |
| **Direction x Hemisphere** | **-0.45** | **0.13** | **< .001\*\*\*** |
| Orientation x PICTS | 0.07 | 0.07 | .33 |
| **Orientation x Infant Carrying** | **-0.15** | **0.07** | **.03\*** |
| **Orientation x STQ-r** | **-0.23** | **0.07** | **.002\*\*** |

The effect of Direction within each Hemisphere was tested using estimated marginal means (Kenward-Roger degrees of freedom). In the right hemisphere, there was no significant difference in neural tracking in the two different movement directions (t(832) = 0.89, p =.37). However, in the left hemisphere, neural tracking was stronger when the movement was in the rightward direction (t(832) = 3.99, p <.001). Importantly, adding the Orientation × Hemisphere interaction or the three-way Orientation × Hemisphere × Direction interaction did not improve model fit (Table 2), indicating that the Hemisphere × Direction effect did not depend on whether the point-light walkers were upright or inverted. Additionally, as adding a three-way interaction with age to the model did not improve model fit either (Table 2.), stronger processing of rightward movement in the left hemisphere appears to be age-independent. The Hemisphere x Direction interaction effect on baseline-subtracted amplitudes is shown in Figure 4.

**Figure 4.**
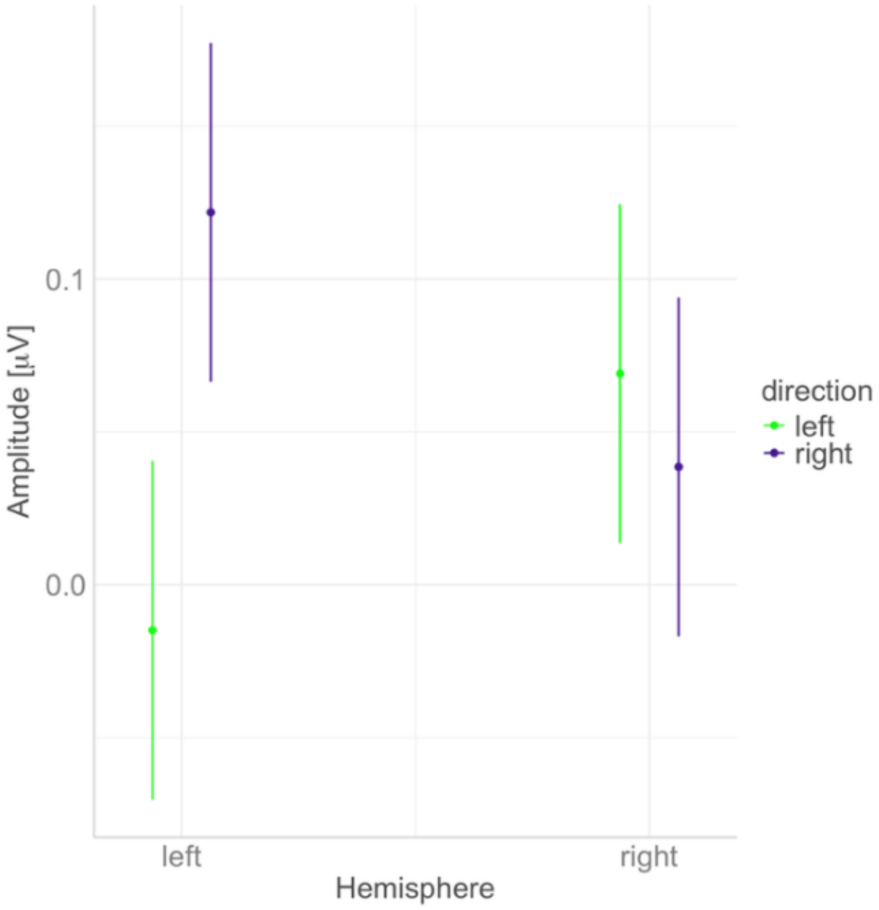
Interaction effect between the hemisphere in which neural tracking is measured and the direction of the point light walkers’ movement on baseline-subtracted amplitudes.

## 4. Discussion

The present study examined neural tracking of visual biological motion rhythms in 3- and 6-month-old infants. We presented infants with an adapted rhythmic visual stimulation paradigm previously applied with adults (Cracco et al., 2022), in which point light walkers were moving at a rhythmic pace of 2.4 Hz while we measured their electrical brain activity using EEG. There were four different conditions: upright leftward movement, upright rightward movement, inverted leftward movement and inverted rightward movement. Neural tracking was quantified as baseline-subtracted amplitudes at the frequency of movement and its first harmonic.

Additionally, we investigated caregiver touch-related practices and attitudes, with a special focus on infant carrying, as possible factors involved in shaping infant neural tracking of biological motion.

First, we hypothesised that infant neural tracking would be stronger for upright, as compared with inverted, biological motion, and that this effect would be more pronounced in older (6 months) than in younger (3 months) infants. However, we found that neural tracking of biological motion rhythms was equally strong for upright and inverted stimuli, and that it did not differ between the age groups. While infant looking behaviour preference for upright, as compared with inverted, biological motion appears to be present from around 2 months of age (Fox & McDaniel, 1982; Sifre et al., 2018), paradigms used to show this preference involved the presentation of both types of stimuli side by side. Studies more closely resembling our procedure, where stimuli were presented sequentially, showed longer looking toward the upright (vs. inverted) biological motion stimuli in newborns (Simion et al., 2008), but not in 6-month-olds (Kuhlmeier et al., 2010) – this is consistent with our findings, as we did not find effects of stimulus orientation on looking times. Further, research examining neural processing of biological motion in 8-month-olds showed orientation condition differences in neural responses between 200 and 300 ms after stimulus onset, measured with event-related potentials to stimuli presented for 1 second (Reid et al., 2006). In contrast, in a functional near infrared spectroscopy study that involved longer presentation periods of several seconds, 8-month-olds’ neural processing of upright and inverted biological motion stimuli was similar between conditions, with both types of stimuli eliciting significant activations in the right superior temporal sulcus area (Lisboa et al., 2020). It is possible that the effects of orientation (upright vs. inverted) may be more evident in earlier stages of visual processing, and in paradigms that measure preferential looking during simultaneous stimuli presentation. Notably, we focused on a rhythm-based measure of biological motion processing, and the visual rhythm of the walking pace was salient in both the upright and inverted conditions. This suggests that infants’ neural tracking of biological motion rhythms, different from adults (Cracco et al., 2022), is less specialized for upright human walking motion. As inversion is thought to perturb configural visual processing of the human body, infants at 3 and 6 months of age don’t seem to rely on global configural processing when tracking visual motion rhythms.

Second, we hypothesised that measures of caregiver-reported practice of infant carrying and other touching behaviours, as well as caregiver attitudes towards social touch, would predict infant neural tracking of biological motion rhythms. We found some evidence for this hypothesis. Caregiver-reported infant carrying positively predicted infants’ neural tracking of biological motion rhythms. More positive caregiver attitudes and affect toward social touch were also linked with stronger neural tracking of biological motion rhythms. Interestingly, for both these effects we found interactions with the orientation of point light walkers, such that both infant carrying as well as caregiver touch attitudes were more strongly positively predictive of neural tracking of inverted, as compared with upright stimuli. No associations with a general measure of infant-directed touching behaviours (including stroking, cuddling, kissing, etc.) were found. Given prior research suggesting that in infants, both upright and inverted biological motion stimuli engage social brain areas involved in biological motion perception in adults (Lisboa et al., 2020), but upright biological motion tends to more readily attract attention (Fox & McDaniel, 1982; Sifre et al., 2018), we speculate that the practice of infant carrying may promote infant recognition of inverted biological motion as a socially salient stimulus. Further, more touch and proximity from the caregiver, as potentially indicated by more positive self-reported attitudes toward social touch in general (Brzozowska et al., 2021), may have similar effects.

Similarly to how touch promotes infant processing of faces with averted gaze (Della Longa et al., 2019) – a “less social” stimulus, as compared with faces with direct gaze – it may promote infant sensitivity to the inverted biological motion stimulus as a potentially socially relevant signal.

Infant familiarity with the rhythms of locomotion, through passively experienced caregiver locomotion while being carried, may further strengthen this effect, especially for the processing of rhythmic biological motion.

This speculative interpretation will have to be more closely examined in future studies, in particular with regards to the causality of the investigated effects. Would experimentally manipulating infant experience with being carried and other types of touch stimulation affect their neural processing of rhythmic biological motion? If there is a causal effect, do social touch and proximity affect infant processing of biological motion through their broad effects on social salience (Della Longa et al., 2019), or is the rhythmic multimodal stimulation associated with being carried (Rocha et al., 2021b) a crucial element, affecting infant neural tracking of biological motion rhythms cross-modally? Finally, using a scrambled vs. intact contrast instead of the upright vs. inverted contrast could yield more insights into the specificity of the effects – we would expect that no associations with neural tracking of scrambled biological motion rhythms would be present, speaking for the specifically social nature of these putative effects. Hopefully, future research will address these outstanding questions.

Lastly, we conducted exploratory analyses that included additional variables such as the direction of the biological motion (left or right) as well as the hemisphere in which neural tracking was measured (left or right). Similarly to gaze direction, infants use biological motion direction as an attentional cue (Bardi et al., 2015; Lunghi et al., 2019). Further, prior research reported right lateralisation of the neural response to biological motion in both adults (Cracco et al., 2022; E. Grossman et al., 2000) and infants (Hirai & Hiraki, 2005; Reid et al., 2006). In our study, while no main effect of hemisphere was found, there was a main effect of movement direction, such that infant brains tracked rightward movement more strongly than leftward movement. Interestingly, an interaction with the hemisphere emerged, whereby the processing of rightward movement was significantly stronger than the processing of leftward movement in the left hemisphere in particular. A study using a looking behaviour-based habituation paradigm reported discrimination between rightward and leftward movement of point light walkers in 6-month-olds only in the upward orientation condition (Kuhlmeier et al., 2010). Similarly, only upright, but not inverted, point light walkers cue 6-month-olds’ visual attention toward the left or right side of a screen, depending on their walking direction (Bardi et al., 2015; Lunghi et al., 2019). In contrast, we found no interaction effects between direction and orientation on neural tracking. While it is difficult to identify what led to the lack of orientation and direction interaction in our study, given the differences between prior research and our study in terms of the paradigm as well as the age of participants, our incidental finding seems to point to interesting lateralisation effects in direction discrimination of biological motion, which future research could shed more light on.

The fact that we observed robust neural tracking of visual rhythms of biological motion in 3-and 6-month-olds is promising with regards to possible future applications of this approach based on rhythmic visual stimulation. Future directions include research examining links between the strength of neural tracking of biological motion rhythms in early infancy and the development of abilities that rely on biological motion processing, such as imitation (Happé et al., 2017). Additionally, as scientific interest in the role of rhythmicity in early caregiver-infant interactions is increasing (Alviar et al., 2025), it is crucial to understand how the processing of rhythmicity of relevant visual rhythms develops in infancy. In particular, neural tracking of interactional and communicative rhythms is likely at the core of early abilities to synchronise with others behaviourally and neurally (Hoehl et al., 2025). As researchers continue to explore the role of rhythm processing in early neurocognitive development, our study contributes evidence focused specifically on biological motion rhythms.

