## Supplementary Materials for "Neural tracking of biological motion rhythms in early infancy: links to caregiver touch-related behaviours and attitudes"

Supplementary materials accompanying the manuscript titled "Neural tracking of biological motion rhythms in early infancy: links to caregiver touch-related attitudes and behaviours"

S1. Minor deviations from preregistration

The preregistration document for the study can be found under this link: https://aspredicted.org/ev22y6.pdf

Minor deviations from the preregistered analysis were as follows:

a) Instead of segmenting the data into 1-second-long segments, we segmented them into 1.668-second segments. This led to the segments being time-locked to the beginning of the walking cycle, which, in turn, allowed us to average the segments before calculating the power spectrum (rather than averaging power spectra), increasing signal to noise ratio (Benjamin et al., 2021).
b) Instead of conducting linear regression analyses to address the two preregistered hypotheses, we analyzed the data with mixed-effects models. Mixed-effects models were considered preferable to linear regression models as they better account for our data structure and mitigate statistical power loss due to data missingness (Muradoglu et al., 2023).
c) Due to the 7.2 Hz harmonic not reaching significance (see main manuscript text, section 3.2), we only included baseline-subtracted amplitudes at 2.4 Hz and 4.8 Hz in the analysis.

In other aspects, we closely followed the preregistration.

S2. Reasons for data missingness

| Age group | Number of infants who participated | Number of valid datasets | Reasons for missingness |
|---|---|---|---|
| 3 months | 41 | 31 | no EEG collected due to fussiness (2); no photodiode data/experimenter error (2); cross-correlation value across all frequencies before and after wavelet |

| | | | |
|---|---|---|---|
| | | | thresholding below 0.1 (4); data deemed too noisy upon visual inspection after preprocessing (2) |
| 6 months | 41 | 30 | no photodiode data/experimenter error (1); cross-correlation value across all frequencies before and after wavelet thresholding below 0.1 (8); data deemed too noisy upon visual inspection after preprocessing (2) |
